# MecCog: A knowledge representation framework for genetic disease mechanism

**DOI:** 10.1101/2020.09.03.282012

**Authors:** Kunal Kundu, Lindley Darden, John Moult

## Abstract

**Motivation:** Experimental findings on genetic disease mechanisms are scattered throughout the literature and represented in many ways, including unstructured text, cartoons, pathway diagrams, and network graphs. Integration and structuring of such mechanistic information will greatly enhance its utility.

**Results:** MecCog is a graphical framework for building integrated representations (mechanism schemas) of mechanisms by which a genetic variant causes a disease phenotype. A MecCog mechanism schema displays the propagation of system perturbations across stages of biological organization, using graphical notations to symbolize perturbed entities and activities, hyperlinked evidence tagging, a mechanism ontology, and depiction of knowledge gaps, ambiguities, and uncertainties. The web platform enables a user to construct, store, publish, browse, query, and comment on schemas. MecCog facilitates the identification of potential biomarkers, therapeutic intervention sites, and critical future experiments.

**Availability and Implementation:** The MecCog framework is freely available at http://www.meccog.org.

**Supplementary information:** Supplementary material is available at *Bioinformatics* online.

## INTRODUCTION

Findings from experimental studies of disease mechanism are reported across multiple publications in varying combinations of structured and unstructured data and many different diagrammatic representations. A number of projects have addressed different aspects of the resulting knowledge integration problem. These include building disease-specific knowledge management resources (for example alzforum.org (Kinoshita and Clark, 2007)) and ontologies (ADO (Malhotra *et al*., 2014), PDON (Younesi *et al*., 2015), CVDO https://bioportal.bioontology.org/ontologies/CVDO); compiling disease etiology databases (HGMD (Stenson *et al*., 2017), ClinVar (Landrum *et al*., 2018), CIVIC (Griffith *et al*., 2017), PanelApp (Martin *et al*., 2019)); development of biomedical text mining methods (DARPA’s Big Mechanism program (Cohen, 2015)); development of statistical methods for evidence integration and assessment (Konopka and Smedley, 2020); and community-driven expert systems medicine disease maps projects (Mazein *et al*., 2018). Each of these contributes elements of a solution, but a major omission is an integrated representation of mechanism knowledge in a clear, precise, and comprehensive manner.

There have also been major technological advances in the development of tools to support mechanism descriptions, such as graphical notations (SGBN (Systems Biology Graphical Notation) (Novère *et al*., 2009)) and languages (SBML (Systems Biology Markup Language) (Hucka *et al*., 2018), KGML (KEGG Markup language - https://www.genome.jp/kegg/xml/), BCML (Biological Connection Markup Language) (Beltrame *et al*., 2011), BioPAX (http://www.biopax.org/), BEL (Biological Expression Language - https://bel.bio/)) to encode representations; software to draw and visualize models (GO-CAM (Thomas *et al*., 2019), PathWhiz (Pon *et al*., 2015), Cytoscape (Shannon *et al*., 2003)); linked data formats such as Nanopublications (Mina *et al*., 2015) to organize provenance and metadata for scientific assertions; and databases to store and query graph-based representations (Neo4j - https://neo4j.com/, (Himmelstein *et al*., 2017)).

With the help of these tools; pathway, network, and disease map representation types have been created to describe aspects of biological system mechanism and in some cases disease mechanisms as well. For instance, KEGG (Kanehisa *et al*., 2016), and Reactome (Fabregat *et al*., 2017) pathways represent normal and perturbed molecular interactions that are part of cellular or metabolic processes. STRING (Szklarczyk *et al*., 2018) and GeneMANIA (Franz *et al*., 2018) networks represent integrated information on protein-protein interactions and associations that are part of normally functioning biological systems. Gene ontology (GO) causal activity models (Thomas *et al*., 2019) integrate GO annotations to generate larger models of normal biological function (such as ‘pathways’) in a semantically structured manner. The Disease Maps Project (Mazein *et al*., 2018) provides an encyclopedic description of disease-related signaling, metabolic, and gene regulatory processes. Although together these representations aptly describe the normal working of biological systems, representation of the disease related perturbations is limited. In the existing representations (such as KEGG or Reactome disease pathways), disease state perturbations and consequences are added locally to the depiction of the normal state of the biological system. Adding disease perturbation information to already complex pathway diagrams can be useful, but limits clarity. Also, uncertainties, ambiguities, and ignorance in mechanistic knowledge are not presented in most representations (with the exception of Reactome pathway diagrams, but these label uncertain reaction types only). Such knowledge gaps exist in almost all disease mechanisms (Greenberg and Amato, 2004; Kametani and Hasegawa, 2018).

These considerations led us to propose a graphical framework with an integrated representation of genetic disease mechanisms from gene to phenotype. Our design goals were that the representation framework depict mechanism components across stages of biological organization; display perturbation propagation; make use of standard biomedical ontology terms wherever possible to name the components; provide an intuitive way to visualize ignorance, uncertainties, and ambiguities; and allow tight linkage to evidence in the literature and databases. The MecCog mechanism representation framework (Darden *et al*., 2018) incorporates all these features. The representation formalism is based on the analysis of biological mechanisms developed in the philosophy of biology (Craver and Darden, 2013): Mechanisms are characterized as entities and activities organized such that they are productive of regular changes from start or set-up to finish or termination conditions. In MecCog, a mechanism by which a genetic variant causes a disease phenotype is represented as a *mechanism schema* that displays the propagation of entity and activity perturbations across biological organizational stages (DNA→RNA→Protein→Complex→Organelle→Cell→Tissue→Organ→Phenotype) in the form of a graph (nodes are biological entities; directed edges are causal and labeled with productive activities) constructed from information in the biomedical literature in addition to established biological concepts. The schema structure uses graph properties such as branching, merging, and looping of sub-paths.

In this article, we describe the implementation of the MecCog framework as a web platform with a collaborative environment to manually construct, store, publish, browse, query, and comment on mechanism schemas for genetic diseases. The schema building tool in MecCog is supported by specially designed graphical notations, curated ontology-informed terminology for the annotation of mechanism components (entities and activities), an interactive graphical user interface (GUI) to construct the schema drawings, application programming interfaces (APIs) to fetch reference information and scientific figures, tight integration and hyperlinking of evidence to the graphics, and a secure server to save schemas as JSON (JavaScript Object Notation) objects. The platform supports edit, version, and share operations on each schema to facilitate collaborative work. Mature schemas can be published on the platform, thereby adding to the collection of disease mechanisms available for browsing by MecCog web-site visitors. Sketchier schemas with gaps, ambiguities, and uncertainties can also be published to indicate where additional work needs to be directed.

## METHODS AND RESULTS

### Mechanism schema representation structure

In MecCog, a mechanism by which a genetic variant causes a disease phenotype is represented by multiple steps. Each step consists of a triplet with an input substate perturbation, a mechanism module, and an output substate perturbation (SSP-MM-SSP). A substate perturbation represents a perturbed biological entity (e.g. a DNA base change, altered stability of a protein, altered abundance of a molecular complex, altered state of a cell). A mechanism module represents the productive activity (e.g. transcription, translation, or protein-protein interaction) by which the input sub-state perturbation produces the output sub-state perturbation. The succession of overlapping SSP-MM-SSP triplets represents perturbation propagation across stages of biological organization (DNA, RNA, Protein, Complex, Organelle, Cell, Tissue, Organ, and Phenotype), and together form a mechanism schema. In MecCog, a schema is represented as a graph where the nodes are SSPs and edge labels are MMs, as illustrated in Figure 1.

**Figure 1.**
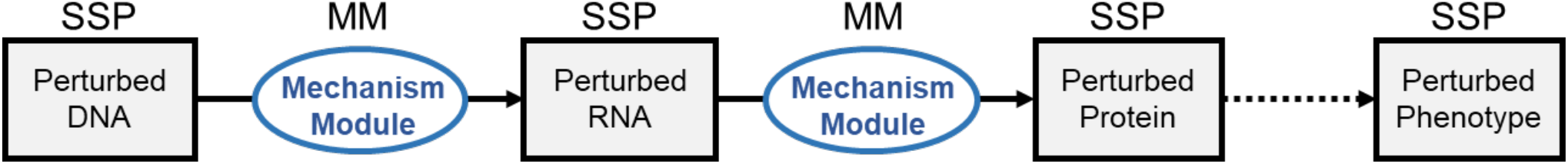
Principles of a mechanism schema. SSP: Substate Perturbation; MM: Mechanism Module. Each SSP represents a perturbed biological entity and each MM represents a productive activity (or a group of entities and activities) that produce an output SSP.

Evidence about SSPs and MMs is curated from the literature. Ambiguities in a mechanism and possible alternative mechanisms are represented in a schema by branching. Branch points may be labeled with the logical operators AND, OR, or AND/OR. The degree of confidence as to whether each SSP and MM is part of a schema is indicated by an evidence strength color code (red least confidence to green most confident) for the corresponding symbol. In addition to SSPs and MMs, five other types of mechanism components are defined in MecCog: 1. *Unknown mechanism module* to represent ignorance about a mechanism component; 2. *Biomarker* to represent entities correlated with a disease phenotype; 3. *Environmental factor* to represent relevant external factors; 4. *Hypothetical therapeutic intervention site*; 5. *Known therapeutic intervention site*.

### MecCog platform web-architecture

Figure 2 shows the web-architecture of MecCog. On the server-side, Node.js (an open-source JavaScript runtime environment) is used as the web-server, Sails.js is used to build the model-view-controller compliant web-application, and a MySQL relational database is used to store data on users and mechanism schemas. The MySQL database is connected to the web-application by the Object-relational mapper (ORM), Waterline, in Sails and all the database transactions use REST APIs secured by CSRF (Cross-site request forgery). The front-end GUI of MecCog is implemented using HTML, CSS, and Javascript, and is made responsive by Bootstrap.js javascript library. The schema building and visualizing GUI is powered by the Rappid Diagramming Javascript library (https://www.jointjs.com/). Rappid also provides a feature for converting diagrams to JSON format and for communicating with the database via AJAX requests. An open-source version of the IntenseDebate commenting system (https://www.intensedebate.com/) is used to render commenting forms.

**Figure 2.**
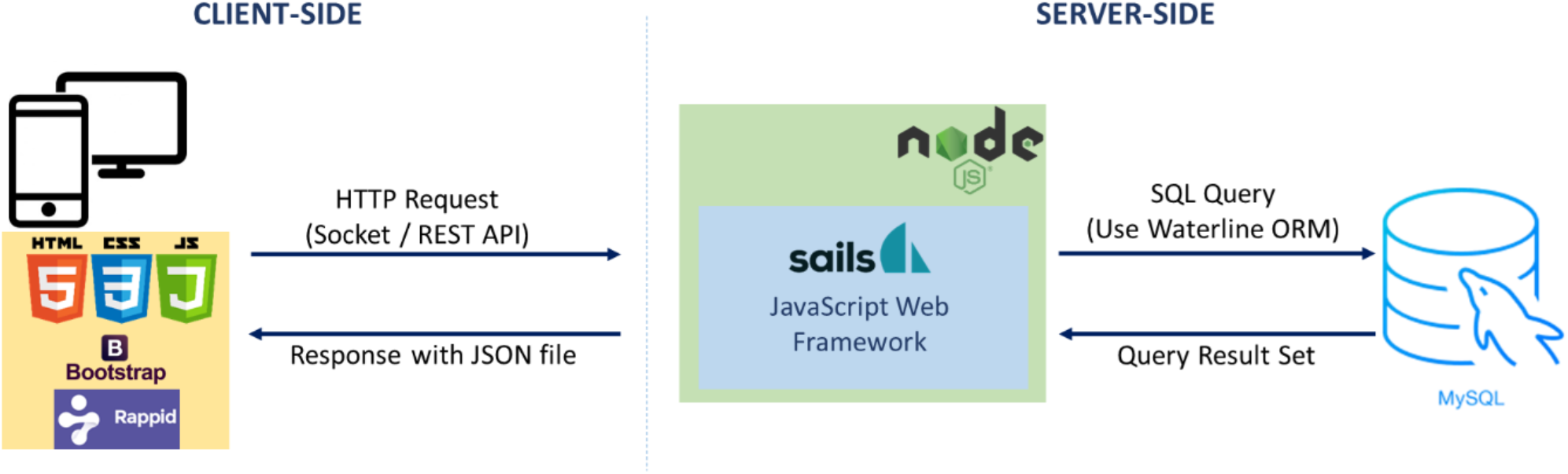
MecCog Web-Application Architecture. HTTP=Hypertext Transfer Protocol, REST API = Representational State Transfer Application Programming Interface, JSON=JavaScript Object Notation, SQL = Structured Query Language, ORM= Object-relational mapping.

### Graphical notation of mechanism components in MecCog

Graphical notations symbolize components of a mechanism schema (Figure 3). An SSP (substate perturbation) is represented by a rectangle containing three types of information – the biological stage where the SSP occurs, the perturbation class name (curated from standard biomedical ontologies wherever possible), and the instance of that perturbation class. For example, a truncated *NOD2* protein can be represented by an SSP with *Protein* as the stage name, *Truncated Protein* as the SSP perturbation class name (from BioAssay Ontology (Visser *et al*., 2011)), and *NOD2:1007fs* as the instance of the SSP class. A biomarker is a special case of an SSP and is represented by the same shape but with a different color. An environmental factor is represented by a cloud icon. Known and hypothetical therapeutic intervention sites are represented by pink and blue octagons respectively. A known mechanism module is represented by a clear oval displaying the MM class or instance name, such as transcription or protein folding. An unknown mechanism module is represented by a black oval.

**Figure 3.**
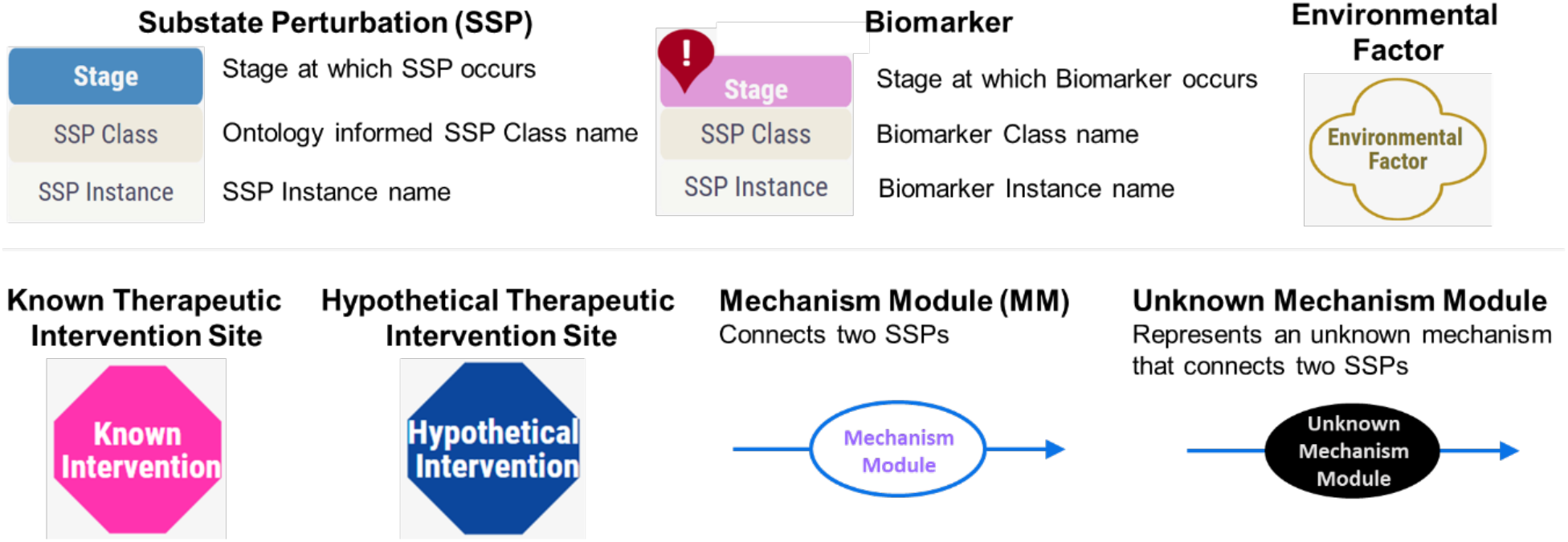
Graphical notations for components in a mechanism schema.

### Mechanism schema meta-information and schema component annotations in MecCog

Table 1 summarizes the mechanism schema data model. Table 1A shows the meta-information of a schema. Each schema in MecCog is identified by a unique accession number automatically generated by the platform. Schema authors provide a schema name, a schema caption, genes that are part of the schema, keywords relevant to the mechanism, names of authors who constructed the schema, and the name of the curator who publishes the schema, monitors comments, and approves changes. Authors also provide a schema description with scientific background information.

Table 1B shows the mechanism component annotations. All mechanism components in a schema are annotated with a unique component identifier. Nine stage names may be assigned to an SSP component notation – DNA, RNA, Protein, Complex, Organelle, Cell, Tissue, Organ, and Phenotype. For the molecular stages (DNA, RNA Protein, and Complex), a set of stage-specific SSP perturbation class names, such as SNV, mRNA abundance, or protein stability, have been compiled. Molecular stage MM classes, such as transcription, translation, and protein folding, are also defined (Table 2). Whenever possible the SSP and MM class names are curated from existing biomedical ontologies. Currently used ontologies are listed in Table 2. Where required, ontology terms may be prefixed with a modifier – increased, decreased, or altered. A MecCog schema builder may choose from the curated set of classes for a step, or may add new class names if needed. SSP and MM instance names are in free text. We are in the process of developing a disease mechanism ontology, based on the class names. Such an ontology is potentially useful for automatic text mining of SSP, MM, and triplet information from the literature, so speeding schema building by identifying relevant papers and sections of papers. Environmental factor names are in free text. Therapeutic intervention site components may be annotated with a potential therapy name or known drug name.

**Table 1.**
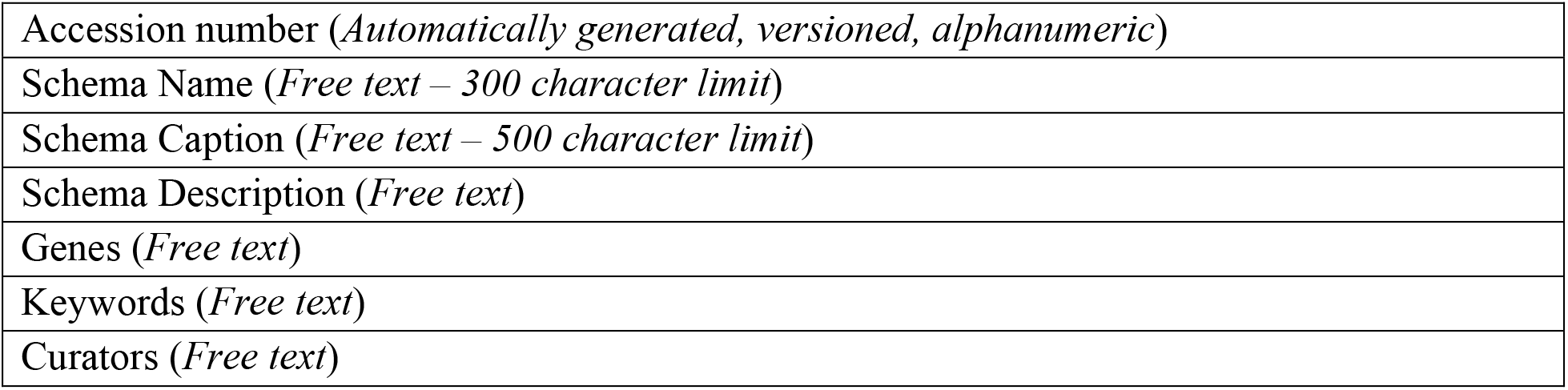
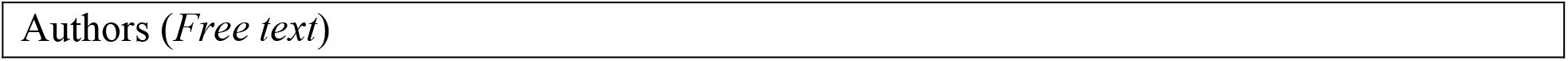
Data Model of mechanism schema and component annotations. Text in parentheses indicates the data type. Table 1A. Mechanism schema meta-information

**Table 1B.**
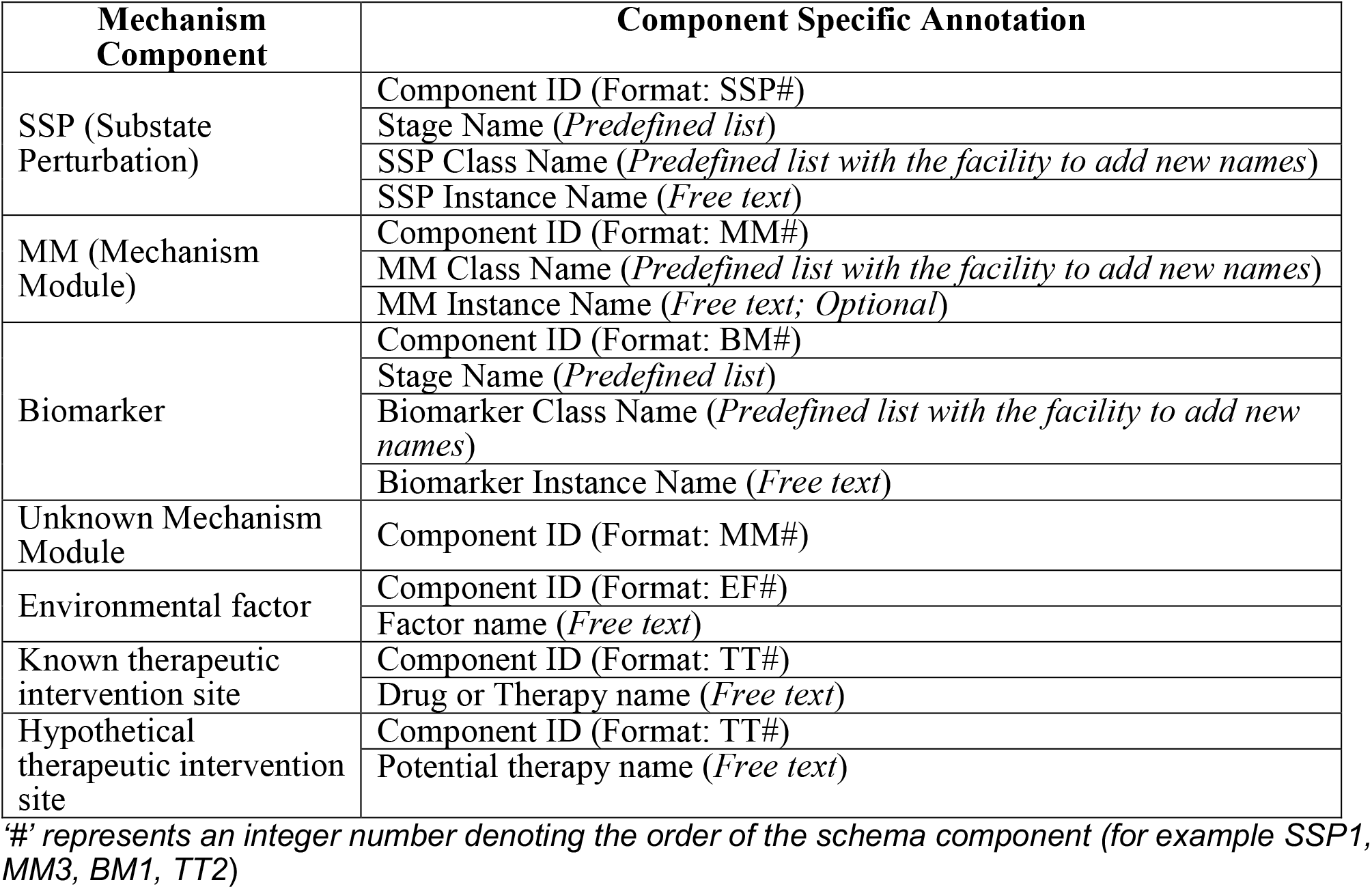
Mechanism component annotations

| Mechanism Component | Component Specific Annotation |
| --- | --- |
| SSP (Substate Perturbation) | Component ID (Format: SSP#) |
|  | Stage Name (*Predefined list*) |
|  | SSP Class Name (*Predefined list with the facility to add new names*) |
|  | SSP Instance Name (*Free text*) |
| MM (Mechanism Module) | Component ID (Format: MM#) |
|  | MM Class Name (*Predefined list with the facility to add new names*) |
|  | MM Instance Name (*Free text; Optional*) |
| Biomarker | Component ID (Format: BM#) |
|  | Stage Name (*Predefined list*) |
|  | Biomarker Class Name (*Predefined list with the facility to add new names*) |
|  | Biomarker Instance Name (*Free text*) |
| Unknown Mechanism Module | Component ID (Format: MM#) |
| Environmental factor | Component ID (Format: EF#) |
|  | Factor name (*Free text*) |
| Known therapeutic intervention site | Component ID (Format: TT#) |
|  | Drug or Therapy name (*Free text*) |
| Hypothetical therapeutic intervention site | Component ID (Format: TT#) |
|  | Potential therapy name (*Free text*) |

#' represents an integer number denoting the order of the schema component (for example SSP1, MM3, BM1, TT2)

**Table 1C.** Evidence annotations for the mechanism components

|  |
| --- |
| For Evidence PubMed IDs (*PMID number*) |
| Against Evidence PubMed IDs (*PMID number*) |
| Figure links from PubMed Central (*PMC figure number*) |
| Confidence score (*Predefined integer score range 1 to 5*) |
| Evidence comment (*Free text*) |

**Table 2.**
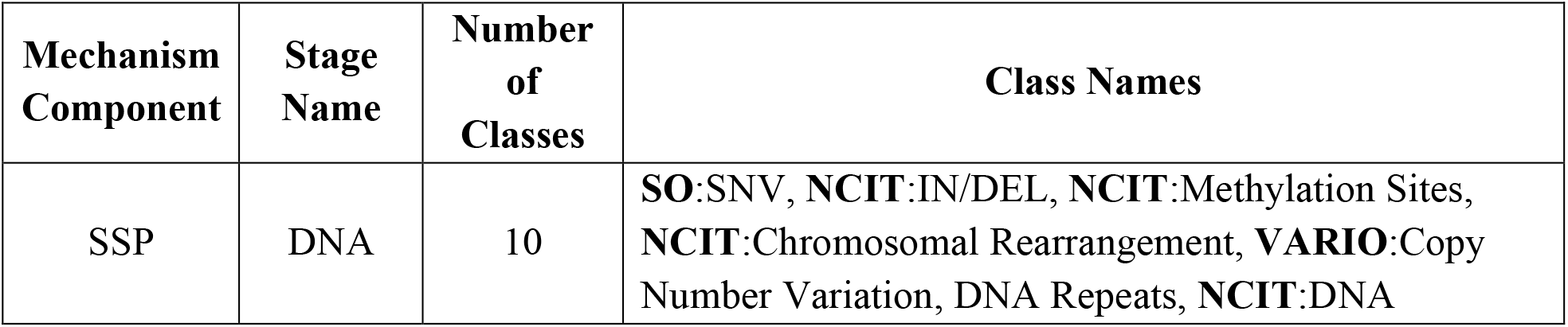
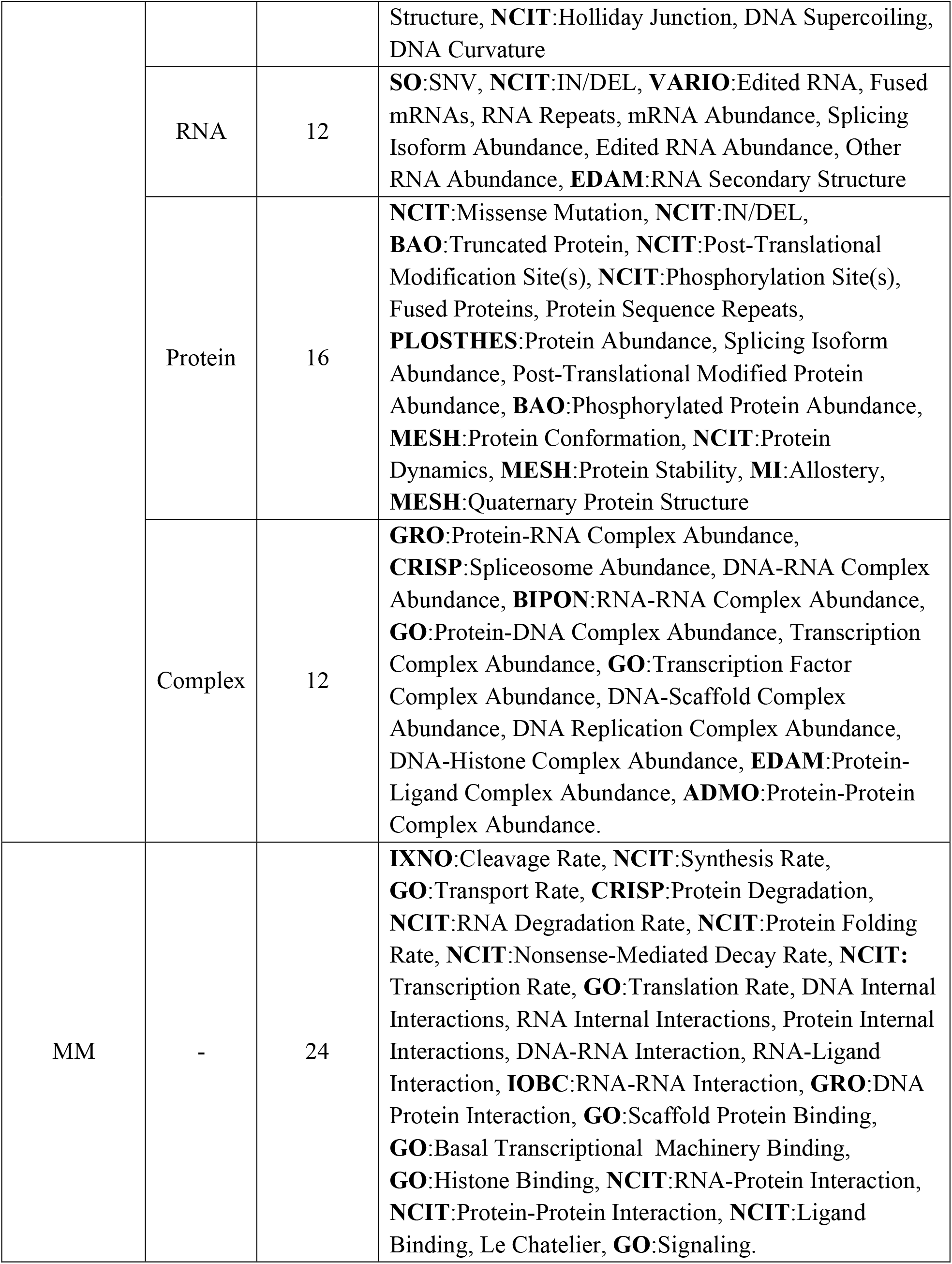
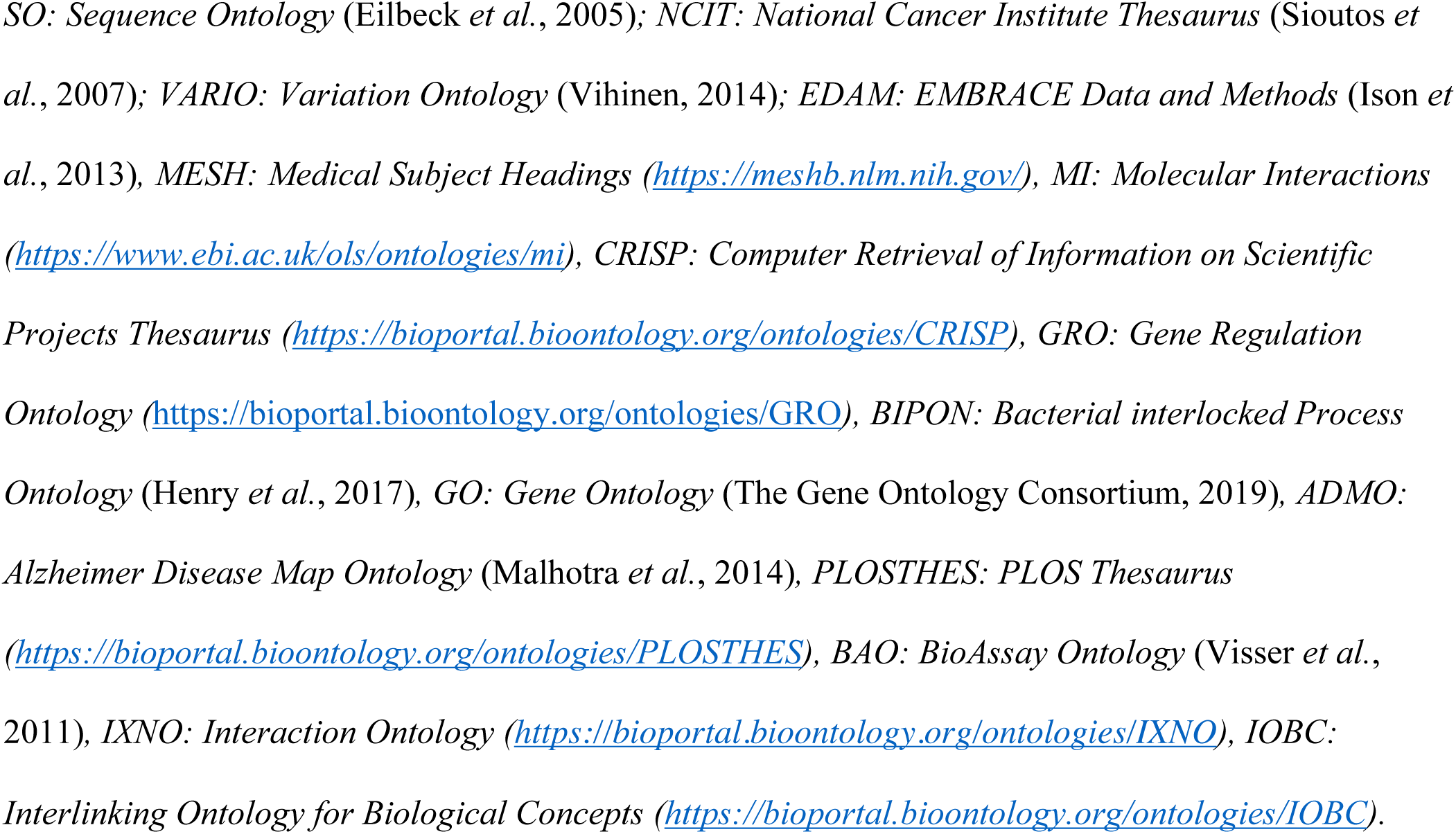
Curated class names for Substate Perturbations (SSP) and Mechanism Modules (MM) at the molecular stages. The class names are curated from biomedical ontologies and are prefixed with the ontology name abbreviation.

For all mechanism components, five types of evidence annotation are defined (Table 1C): 1. *For Evidence PubMed IDs* of papers that contain data supporting a component’s role in a mechanism; 2. *Against Evidence PubMed IDs* for papers that provide data suggesting a mechanism component is incorrect; 3. *Links to figures in PMC* that illustrate aspects of a schema by summarizing experimental results and evidence for spatial and structural features; 4. User assigned *Confidence scores* with five levels (also used to automatically encode a component’s confidence color) based on the strength of the available evidence; 5. *Evidence Comments -* brief free text comments that summarize the evidence.

### Rules for constructing mechanism schemas in MecCog

1. Each schema begins with a genome perturbation and ends in perturbation of a disease-related phenotype such as greater risk of a disease.
2. Overall, the sequence of SSPs in a schema progresses through successive stages of biological organization, from DNA, through RNA, proteins, macromolecular complexes, organelles, cells, tissues, organs, and finally to a phenotype. There may be one or more or no SSP at any particular stage of organization and the order of the stages need not follow a prescribed order. For instance, the schema for Lynch syndrome (http://www.meccog.org/mchain/showpubchain?accession=MS020700047.3), where a causative mutation results in decreased DNA mismatch repair, reverts to the DNA stage after stages involving macromolecular complexes.
3. Each pair of SSPs is linked by an MM. The granularity of an MM may be a single activity (such as splicing, protein-protein interaction, ligand binding, or protein folding) or may represent telescoped combinations of entities and activities (such as protein synthesis, or cell-cell signaling). If an activity is unknown, the black oval unknown mechanism module notation is used.
4. Class names for the SSPs and MMs at the molecular stages (DNA, RNA, Protein, and Complex) can be selected from the pre-compiled list shown in Table 2. If existing names are inadequate, new names can be used. At higher organizational stages, class names are user-provided. Wherever possible these should be part of existing biomedical ontologies. The NCBO BioPortal site (https://bioportal.bioontology.org/) is a source for ontology terms. Since most ontologies describe the normal state of a system, a user may select one of the in-built modifiers (increased, decreased, altered) to prefix a class name so as to represent a perturbed state.
5. An evidence-based confidence score (on the scale of 1 to 5, where one indicates low confidence and five indicates high confidence) should be assigned to each SSP and MM. Evidence on which a confidence score is based should be recorded in the form of supporting/contradicting PMIDs and PMC figure URLs, together with appropriate free text commentary.
6. Two or more possible sub-paths can exist in a schema either because of ambiguity due to conflicting evidence, or alternative sub-mechanisms. Branch points should be labeled with OR, AND or AND/OR.
7. Schemas should explicitly include steps only where there is a perturbation from the normal system. Where the function of a portion of a schema is unperturbed, for example, representing the standard activity of transcription operating on a perturbed input DNA sequence or a standard cell signaling process operates with more or less input signal, that section of the schema should be telescoped into a single mechanism module.

### Steps in constructing, managing, and publishing mechanism schemas

Before beginning schema building a new user must register on the MecCog platform. A registered user may select the “Build Schema” tab to initiate building a new schema or the “My Schemas” tab to access the workspace for managing and editing their existing schemas. Figures 4A and 4B show the two interfaces used in schema construction: A. The *Initiate Mechanism Schema* form used to enter meta-information about a schema, and B. The *Schema Builder* GUI used to draw a schema. In the schema builder, mechanism components can be dragged and dropped from the mechanism component catalog panel to the drawing board panel. Clicking on a component displays five associated control icons: i) Icon to connect to other components; ii) Icon to adjust the component size; iii) Icon to clone the component; iv) Icon to show the pop-up box; and v) Icon to delete the component. Clicking on a component also renders a component-specific annotation form on the rightmost panel of the interface (labeled in Figure 4B). This form is used to enter the stage, class, and instance name of the components, prefix class names with a perturbation type if needed (increased, decreased, or altered), and record the evidence annotations (listed in the Evidence Annotations Table 1C). This is a dynamic form that automatically provides predefined possible perturbation class names for the selected stage (listed in Table 2) and creates fields for adding new PubMed IDs and PMC image URLs. The NCBI E-utilities application programming interface (API) is used on the server-side to fetch publication details for the PubMed IDs. All the evidence annotations are transferred to the current component pop-up box together with hyperlinked PMIDs and PMC image URLs. The pop-up box can be visualized by clicking the *‘i’* shaped control icon of the component. Confidence score values selected in the annotation form are used to automatically apply the appropriate color to the current schema component (red: score 1, orange: scores 2, 3, 4 and, green: score 5). The color of the edge connecting two components is inherited from the target component, so indicating causal confidence. Schemas are saved to the database using the *Click to save* button. For each schema, a unique accession number is automatically generated in the database. The accession number format has a section indicating the version (default is .1) of the schema. The schema builder GUI also has a panel of interactive buttons to undo, redo, clear page, zoom, auto-layout, export (in SVG and PNG formats), and print schema diagrams.

**Figure 4.**
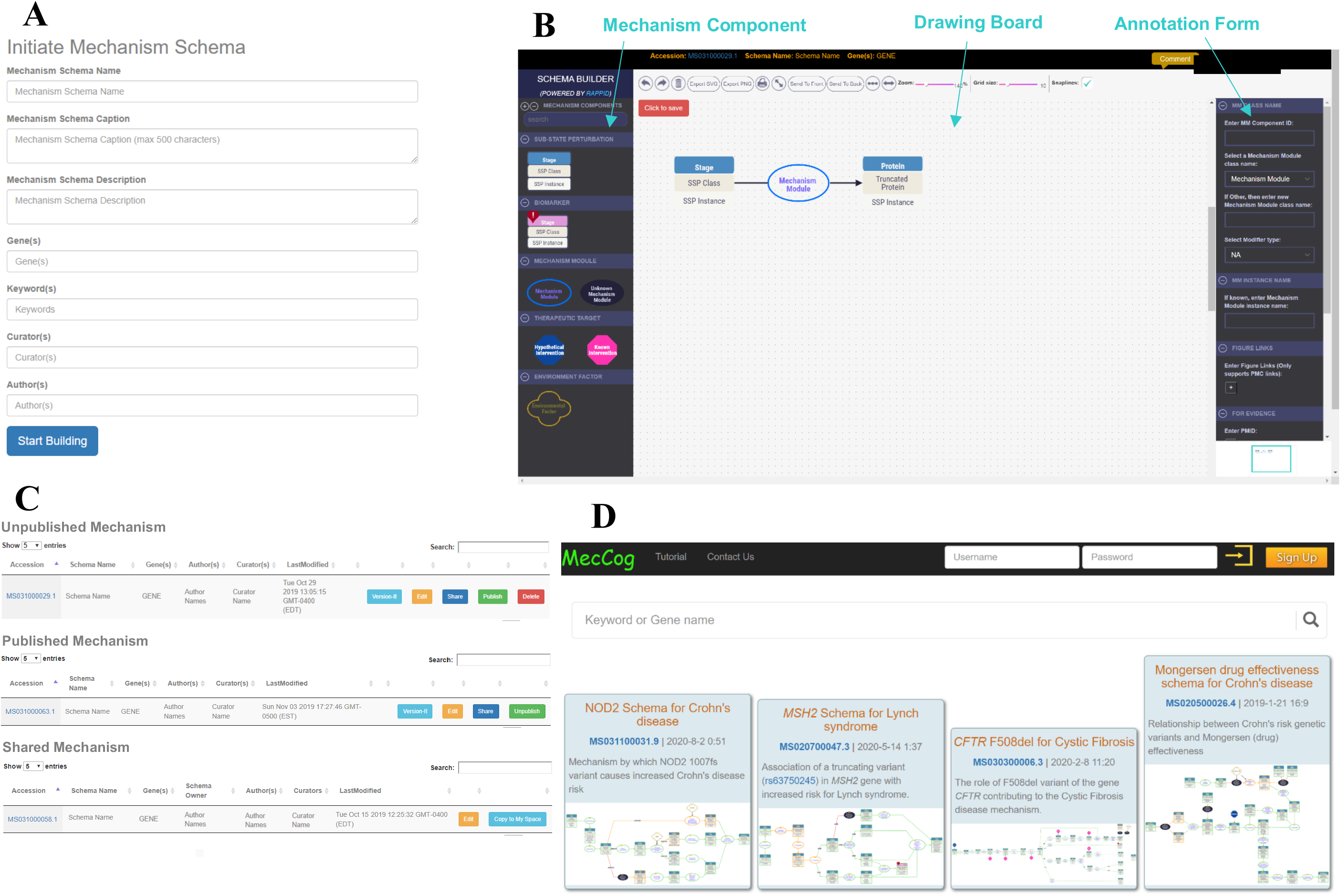
Graphical User Interface (GUI) to construct, manage, and browse mechanism schemas. Figure 4A shows the form for entering meta-information on a schema. Figure 4B shows the schema builder interface, including the mechanism component catalog panel (left), the drawing board panel, and the annotation form panel (right). Figure 4C shows the user workspace interface. Figure 4D shows a portion of the MecCog main webpage, designed to facilitate browsing publicly available mechanism schemas.

Figure 4C shows the view of a registered user’s workspace. Each schema can be versioned, edited, shared with other MecCog users, published, or deleted, using operation-specific buttons. The workspace has three sections: i) The Unpublished Mechanism Schemas section catalogs work-in-progress schemas. The Published Mechanism Schemas section catalogs published schemas. A button to remove each of these from the public collection is provided. iii) The Shared Mechanism Schemas section catalogs schemas that have been shared with the current user. For schemas with edit access privilege, the *Copy to My Space* operation is enabled, allowing the creation of a copy for the user to work on independently. All the schema accession numbers in the workspace page are hyperlinked to the schema specific landing page (described in the next section).

Published schemas are available for browsing via the main webpage of the MecCog site (as shown in Figure 4D) without the need for logging in. There is a search bar that allows schemas to be searched by gene name, keyword, or any component class/instance name. MYSQL’s FULLTEXT indexing (https://dev.mysql.com/doc/refman/5.6/en/innodb-fulltext-index.html) feature is used to support the search operation. The structured organization of mechanistic knowledge in MecCog allows this search to be used to find common entities or activities and common classes used in different schemas. On the main page, schemas are presented in a masonry layout view. Each tile in the view displays the schema name, schema caption, hyperlinked accession number of the schema linking to the corresponding schema landing page (described in the next section), and a hyperlinked schema image linked to an interactive web-based visualization of the schema (also described in the next section).

### Schema landing page, schema visualizer, and schema report

A schema landing page displays schema meta-information in a tabular layout (Figure 5A). A novel feature of this page is the display of references and hyper-linked PubMed IDs providing evidence for each aspect of the schema as well as PMC images selected to illustrate aspects of the mechanism. The *Schema Visualizer* button on the landing page directs a user to the GUI for interactively navigating the mechanism schema (as shown in Figure 5B). The visualizer inherits all the interactive features of the schema builder GUI (described previously). A unique feature of the visualizer is the tight integration of the graphical notations for the mechanism components and the associated evidence information (presented in the pop-up box). The pop-up box (yellow-colored box in Figure 5B) displays hyperlinked ontology sources for the SSP/MM class name (if the term is from an ontology), a brief builder-provided commentary on the evidence, and hyperlinked PMIDs and PMC figure IDs. There is a help icon (‘?’) in the visualizer to display the mechanism schema key. Clicking on the Comment button in the visualizer opens a modal box to view or enter comments about the schema. The *Schema Report* button on the landing page generates a narrative report in which the meta-information, mechanism components, and evidence annotations about the schema are presented in a structured format. The schema content can be downloaded as a JSON file from the landing page using the download icon. The page also has social media share icons.

**Figure 5.**
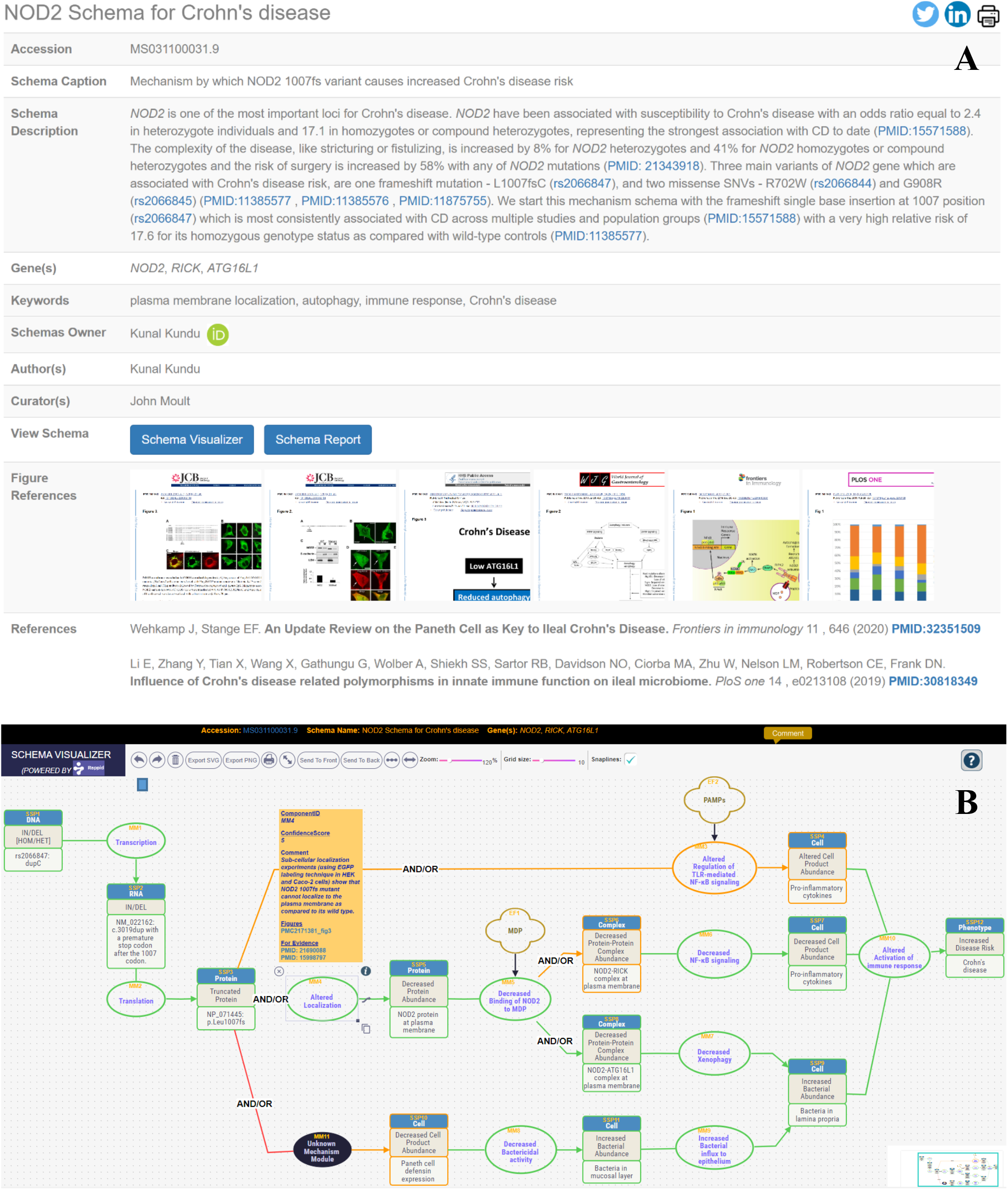
NOD2 mechanism schema entry in MecCog. This schema describes the known mechanisms by which a frameshift mutation (rs2066847) in the NOD2 gene causes an increased risk for Crohn’s disease. Figure 5A shows part of the landing page of the NOD2 schema, displaying the meta-information in tabular format. This page includes the collection of thumbnails of PMC figures selected to illustrate aspects of the mechanism and the list of references with PubMed IDs from which evidence was derived (the list is truncated here - there are 26 references). Figure 5B shows the schema visualizer GUI used for interactive navigation of the schema. For this schema, four possible submechanisms with varying levels of evidence (indicated by the confidence colors – red=low, orange=medium, and green=high) are included. The example yellow pop-up box displays hyperlinked evidence for the associated MM. The comment button on the top can be used to open a modal box, allowing a user to view and add comments. Details of the mechanism are described in the text.

### An example MecCog Schema: Known mechanisms by which a frameshift mutation in the *NOD2* gene causes an increased risk of Crohn’s disease

Figure 5 shows two pages of a MecCog mechanism schema (http://www.meccog.org/mchain/showpubchain?accession=MS031100031.9) describing the mechanism by which a frameshift mutation (rs2066847; NM_022162.3:c.3019dup (p.Leu1007fs)) in the *NOD2* gene causes an increased risk of Crohn’s disease (CD). *NOD2* is the first gene for which variants were found to be associated with altered CD risk (Ogura *et al*., 2001; Hugot *et al*., 2001; Yamamoto and Ma, 2009) and the 1007fs mutation is most consistently associated with CD across multiple studies and population groups (Economou *et al*., 2004) with a very high relative risk of 17.6 for its homozygous genotype status as compared with wild-type controls (Ogura *et al*., 2001). Figure 5A shows the landing page displaying the meta-information of the schema and the list of references used as evidence in the schema, together with figures used to illustrate aspects of the mechanism.

Figure 5B shows the view of the NOD2 1007fs schema in the interactive visualizer. This schema was constructed using information about mechanism reported in 26 research articles. The left-most SSP (SSP1) represents the DNA stage perturbation (i.e. the single base insertion of cytosine - rs2066847 in the *NOD2* gene). The paths in the schema show how the effect of this perturbation propagates through the RNA, protein, complex, and cell stages (represented by the stage-specific SSPs and MMs) so causing the increased Crohn’s disease risk phenotype (SSP12). At the RNA stage (SSP2), the rs2066847 variant causes the insertion of a cytosine after the first nucleotide of codon 1007, so introducing a premature downstream stop codon. This leads to the protein stage perturbation, a truncated NOD2 protein (SSP3) missing the last 33 amino acids of the wild-type sequence (Lécine *et al*., 2007). Following this, the schema branches represent the multiple submechanisms by which the truncated NOD2 protein may lead to the increased Crohn’s disease risk phenotype by altering the activation of the immune response (MM10) (Negroni *et al*., 2018; Strober and Watanabe, 2011; Park *et al*., 2007). All the branches are labeled ‘AND/OR’ since none has fully compelling supporting evidence. The submechanism of each branch is outlined below.

A. The top branch (SSP3→MM3 → SSP4) shows a potential alteration to NOD2-dependent regulation of Toll-like receptor (TLR) mediated NF-κB signaling that produces pro-inflammatory cytokines in response to the pathogen-associated molecular patterns (PAMPs) such as lipopolysaccharide (LPS), or muramyl dipeptide (MDP). This path is sparse and labeled medium confidence (orange) because the mechanism of interaction between NOD2 and TLRs is not known, nor is it clear how that interaction normally results in increased production of pro-inflammatory cytokines (Underhill, 2007). Different models have been proposed to describe the mechanism: synergistic production of TNF-α by NOD2 and TLR4 (Wolfert *et al*., 2002); activation of the inflammasome by NOD2 via RICK to produce IL-1β From pro-IL-1β generated as the result of TLR signaling (Sarkar *et al*., 2006); and MDP (the primary agonist for NOD2 (Grimes *et al*., 2012)) dose-dependent TNF-α production by NOD2 and TLR2 (Borm *et al*., 2008). Further, for none of these possibilities has the effect of the NOD2 100fs variant been investigated. These details are provided in the pop-up box for MM3.
B. The middle branch shows that truncated NOD2 protein has lost its ability to localize to the plasma membrane (MM4 → SSP5) (Barnich *et al*., 2005; Morosky *et al*., 2011) where binding to incoming MDP normally produces an activated state of the protein (Al Nabhani *et al*., 2017). In turn, activated NOD2 forms complexes with RICK and with ATG16L1 (Barnich *et al*., 2005; Travassos *et al*., 2010). The schema shows these effects as lower abundance of the NOD2-RICK complex (MM5 → SSP6) (Barnich *et al*., 2005) and the NOD2-ATG16L1 complex (MM5 → SSP8) (Travassos *et al*., 2010). There is no experimental evidence of the NOD2 1007fs protein’s impact on complex formation. Therefore the MM5 → SSP6 step in the schema is labeled medium confidence (orange). Following this step, the lower abundance of the NOD2-RICK complex alters downstream NF-κB signaling (SSP6→MM6→SSP7) (Caruso *et al*., 2014; Lécine *et al*., 2007; Barnich *et al*., 2005; Girardin *et al*., 2003), resulting in lower pro-inflammatory cytokine production, so contributing to an altered activation of the immune response (MM10) (Vilela *et al*., 2012; Park *et al*., 2007; Strober and Watanabe, 2011; Negroni *et al*., 2018). The perturbation of the NOD2-ATG16L1 complex affects the xenophagy process (autophagy against bacteria) (MM7) (Travassos *et al*., 2010) so leading to an increase in the abundance of bacteria in the lamina propria (SSP9) (Sidiq *et al*., 2016) and thereby likely contributing to a more aggressive response from other components of the immune system, as indicated by the altered activation of immune response (MM10). This sub-path is labeled high confidence (green) as its mechanism components are well understood based on the available evidence in the literature. The yellow pop-up box for MM4 shows an example of an evidence commentary with an associated hyperlinked PMC figure and PMIDs.
C. The lower branch of the schema provides examples of the representation of a gap in knowledge and of overall low confidence. Commensal bacteria are largely prevented from penetrating the gut wall by an outer mucosal barrier and the epithelial cell layer. Paneth cells situated in the gut epithelial layer produce a range of antibiotic defensin peptides to aid in preventing commensal bacteria from traversing the mucosal layer. Some data suggest that this process is partly dependent on MDP binding to NOD2 in these cells, likely signaling that significant numbers of bacteria are getting through to the epithelial cell layer, and so triggering an increased response. Data supporting that view come from an experiment showing stimulation of NOD2 by MDP binding induces production of defensin HNP-1 (human neutrophil peptide 1) in Caco-2 cells (Yamamoto-Furusho *et al*., 2010). It has also been shown that the NOD2 1007fs protein fails to induce the production of defensin hBD2 (human β-defensin-2) in several epithelial cell lines (Voss *et al*., 2006). Hence the link between the presence of the 1007fs variant (SSP3) and increased defensin production (currently SSP10). But the mechanism by which MDP binding to NOD2 normally causes defensin production is unknown, hence the black oval (MM11) linking those two SSPs. There is also evidence from other studies that do not support the mechanism represented by this schema path: In two out of four CD cohort studies (Wehkamp *et al*., 2005; Simms *et al*., 2008; Hayashi *et al*., 2016), and in NOD2 deficient mouse organoids (Wilson *et al*., 2015), the association between NOD2 and defensin was not reproduced. Hence this branch is labeled low confidence (red). Further along this schema branch, the decrease in defensin production (SSP10) leads to an increased abundance of bacteria in the mucosal layer (SSP11) due to decreased bactericidal activity (MM8). In turn, this contributes to increased bacterial abundance in the lamina propria due to increased bacterial influx from the mucosal layer (MM9) and finally leads to the altered activation of the immune response (MM10).

### Representation of biomarker and therapeutic intervention sites in MecCog

Figure 6A shows an example of the use of the biomarker symbol, in part of the Lynch syndrome schema (http://www.meccog.org/mchain/showpubchain?accession=MS020700047.3). In this schema, microsatellite instability is a diagnostic biomarker (Vilar *et al*., 2014) for Lynch syndrome, resulting from defective base mismatch repair machinery, in turn a consequence of a mutation (rs63750245: C>T) in the *MSH2* gene. Figure 6B shows an example of a putative therapeutic intervention site in a Crohn’s disease schema (http://www.meccog.org/mchain/showpubchain?accession=MS020500019.2) describing the mechanism by which a missense variant (rs3197999: G>A; R703C) in the *MST1* gene (coding for Macrophage Stimulating Protein, MSP) increases disease risk. The missense variant causes a lower abundance of the MSP-RON protein complex by one or both of two mechanisms: a weakened protein-protein interaction (Chao *et al*., 2014; Gorlatova *et al*., 2011) and reduced MSP abundance. Lower abundance of the complex is expected to result in reduced intracellular signaling affecting macrophage activation (Wang *et al*., 2002; L. Kretschmann *et al*., 2010; Häuser *et al*., 2012) and/or epithelial cell survival and growth (Danilkovitch *et al*., 2000; Neurath, 2014). An appropriate compound that bridges the structural interface between MSP and RON could restore wild-type abundance of the complex and hence signaling strength and so eliminate the downstream consequences. (Of course, many factors affect whether this is in fact an effective therapeutic strategy.)

**Figure 6.**
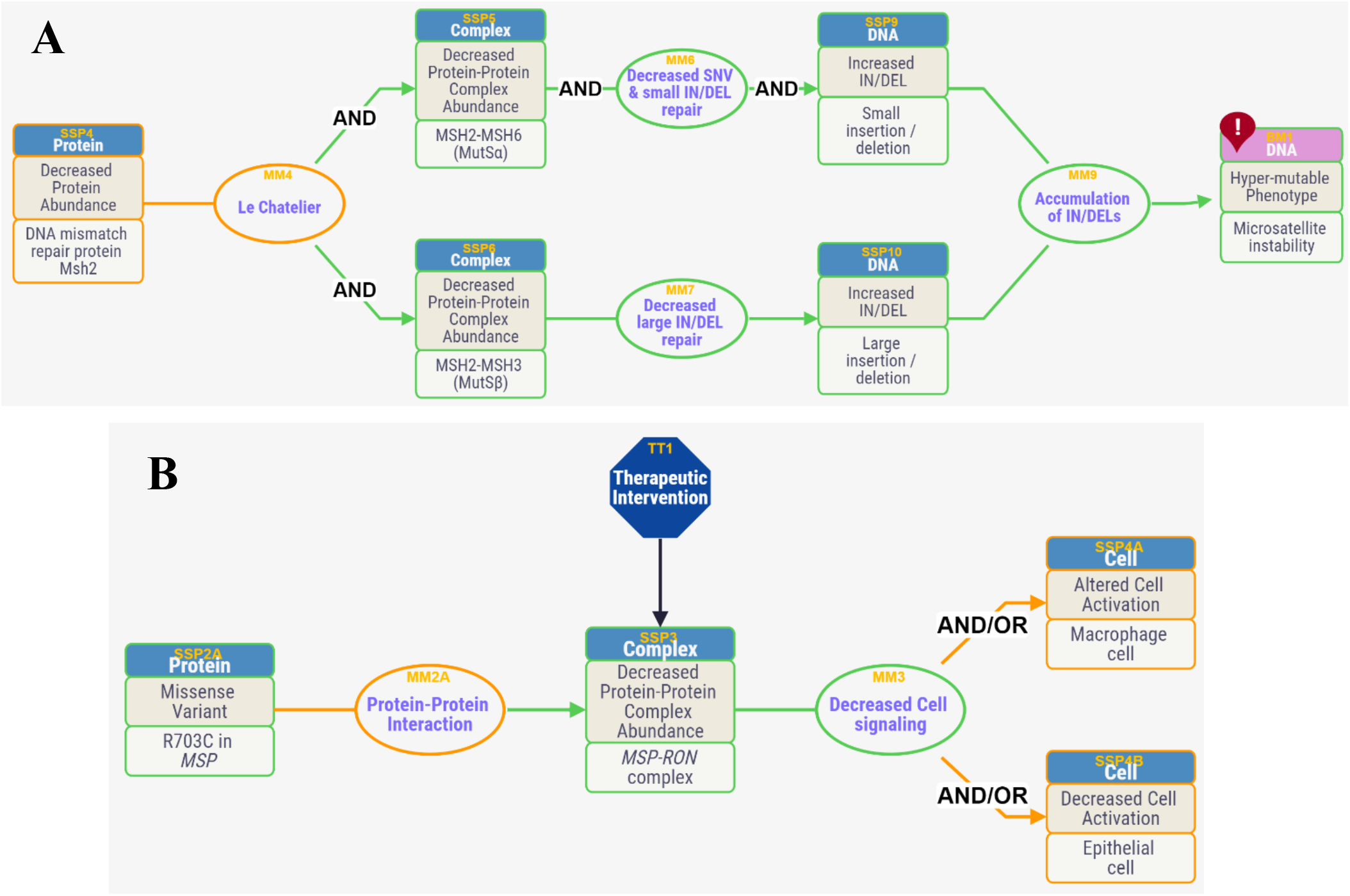
Biomarker and Therapeutic intervention site representation in MecCog. Figure 7A shows part of a Lynch syndrome schema (http://www.meccog.org/mchain/showpubchain?accession=MS020700047.3) where the presence of a nonsense mutation (rs63750245: C>T) in the MSH2 gene causes Microsatellite instability (MSI), a known biomarker (symbolized by the red location icon on the SSP) for the Lynch syndrome. Figure 7B shows part of a Crohn’s disease schema (http://www.meccog.org/mchain/showpubchain?accession=MS020500019.2) where the decreased abundance of the MSP-RON protein complex (SSP3) is a hypothetical therapeutic intervention site, indicated by the blue octagon. In this case, an appropriate small molecule binding across the protein-protein interface might restore the wild-type abundance.

### Validation of the MecCog representation framework

Eleven MecCog mechanism schemas (nine for Crohn’s disease, one for cystic fibrosis, and one for Lynch syndrome) have so far been published, with additional schemas in progress on breast cancer and Alzheimer’s disease. Validation and improvement of MecCog content is obtained by soliciting feedback from specialists in the disease described in each schema. Feedback on the representation technique and platform can be provided using MecCog’s *contact us* form, encouraging users to provide suggestions and report problems. MecCog has also been used as an educational tool for senior undergraduate students in a Human Genetics class at the University of Maryland, providing valuable feedback, for example, linking PMC figures to aspects of schemas.

## Discussion

We have developed MecCog, a graphical knowledge representation framework, to describe genetic disease mechanisms in a structured mechanism schema format. MecCog facilitates the assembly of mechanistic information in terms of perturbation propagation across stages of biological organization, evaluation of the evidence related to that information, and identification of uncertainties, ambiguities, and ignorance. The MecCog web platform provides functionalities to create, store, browse, and search schemas. Graphical notations are annotated with ontology-informed class terms so as to consistently and intuitively represent types of mechanism components found in schemas. The schema interactive visualizer in MecCog tightly integrates the graphics, text, and hyperlinks to evidence sources.

Each schema in MecCog describes mechanisms by which a single genetic variant contributes to the increased risk for the disease phenotype. For complex trait genetic disease and cancer, multiple genetic variants contribute to disease phenotypes (Peter *et al*., 2011; Lilyquist *et al*., 2018). Further, contributions from variants may not be independent, as reflected by evidence of epistatic effects between pairs of variants for complex trait disease (Lin *et al*., 2017; Li *et al*., 2020). The MecCog formalism also supports mechanism schemas with multiple input genetic perturbations. Interactions between these inputs results in a mechanism graph. An example for Crohn’s disease is a barrier integrity mechanism graph constructed by combining schemas on loci relevant to bacterial penetration of the gut-lining mucosal layer (Figure S1). This graph incorporates a number of non-additive interactions between mucin gene variants affecting mucosal-layer integrity (*MUC1, MUC2*), variants affecting the unfolded protein response (*XBP1, ORMDL3*), and variants affecting autophagy (*NOD2, ATG16L1, LRRK2, IRGM*).

Currently, MecCog schemas are manually constructed, relying on human understanding to extract and infer causal connections between mechanism components from literature. Given the scattered and incomplete nature of mechanistic information in literature, this process is complex and requires a combination of prior biological knowledge together with searching for and assimilating new facts and evidence from the literature. These activities are labor-intensive and work best when the schema builder is an expert on the schema subject. To achieve scale for the resource, we require an expert-crowdsourcing strategy, soliciting inputs from appropriate domain experts. The resource is structured so that experts can build schemas based on their knowledge and can also edit and comment on existing schemas. The current version of the MecCog platform supports these activities in the following ways: i) acknowledging contributors to a schema as authors and curators, ii) providing a schema specific commenting interface to solicit input, and iii) allowing versioning of schemas to update content. To implement the crowdsourcing model, we will work closely work with disease-specific research communities (such as IBD Genetics, Crohn’s & Colitis Foundation, and Alzforum).

An obvious question is whether mechanism schemas can be constructed automatically given the structured and unstructured data available in the biomedical domain. The structure of a mechanism schema shares features with that of knowledge graphs (KG), a knowledge representation system initiated by Google in 2012 (https://googleblog.blogspot.com/2012/05/introducing-knowledge-graph-things-not.html). There nodes (aka subjects) represent entities such as real-world objects, events or concepts, and edges (aka predicates) link the nodes with relationships. Because of a KG’s ability to integrate and represent multi-relational databases, many biological KGs (Sosa *et al*., 2019; Celebi *et al*., 2019; Chen *et al*., 2019; Himmelstein *et al*., 2017; PDBe-KB consortium, 2020; https://digitalinsights.qiagen.com/coronavirus-network-explorer/) are being generated, using a combination of manual and automated mining of subject-predicate-object (SPO) triplets from biomedical literature and from bioinformatics relational databases. The elemental SSP-MM-SSP units of a MecCog schema are a subset of SPO triplets and so it is in principle possible to construct a schema by extracting appropriate triplets from a comprehensive knowledge graph. However, preliminary tests of this process suggest that current knowledge graphs do not capture a large fraction of the triplets incorporated in the corresponding mechanism schemas. There are multiple reasons for this, including the absence of biological knowledge in knowledge graphs and the absence of causal reasoning components. We envisage that in the future comprehensive and well-structured KGs will be combined with a repository of biological knowledge and reasoning machines to generate a wide variety of biological mechanisms, as well as providing evaluation of evidence strength and identifying current gaps in mechanism knowledge.

## Acknowledgments

This work was supported in part by the National Institute of Health [R01GM104436 to JM]. KK’s conference travel related to this research was supported in part by NSF award DGE-1632976. We thank Rappid (https://www.jointjs.com/) for providing their JavaScript library under an academic license. We thank Lipika Ray, Yizhou Yin, Maya Zuhl, Christian Presley, and undergraduate students in the University of Maryland Human Genetic course for many useful suggestions and comments on MecCog software. We thank Mark Tonelli for feedback on the MecCog framework and for reviewing the cystic fibrosis schema.

