## Supplementary material for "MecCog: A knowledge representation framework for genetic disease mechanism": Figure S1

Supplementary Figure

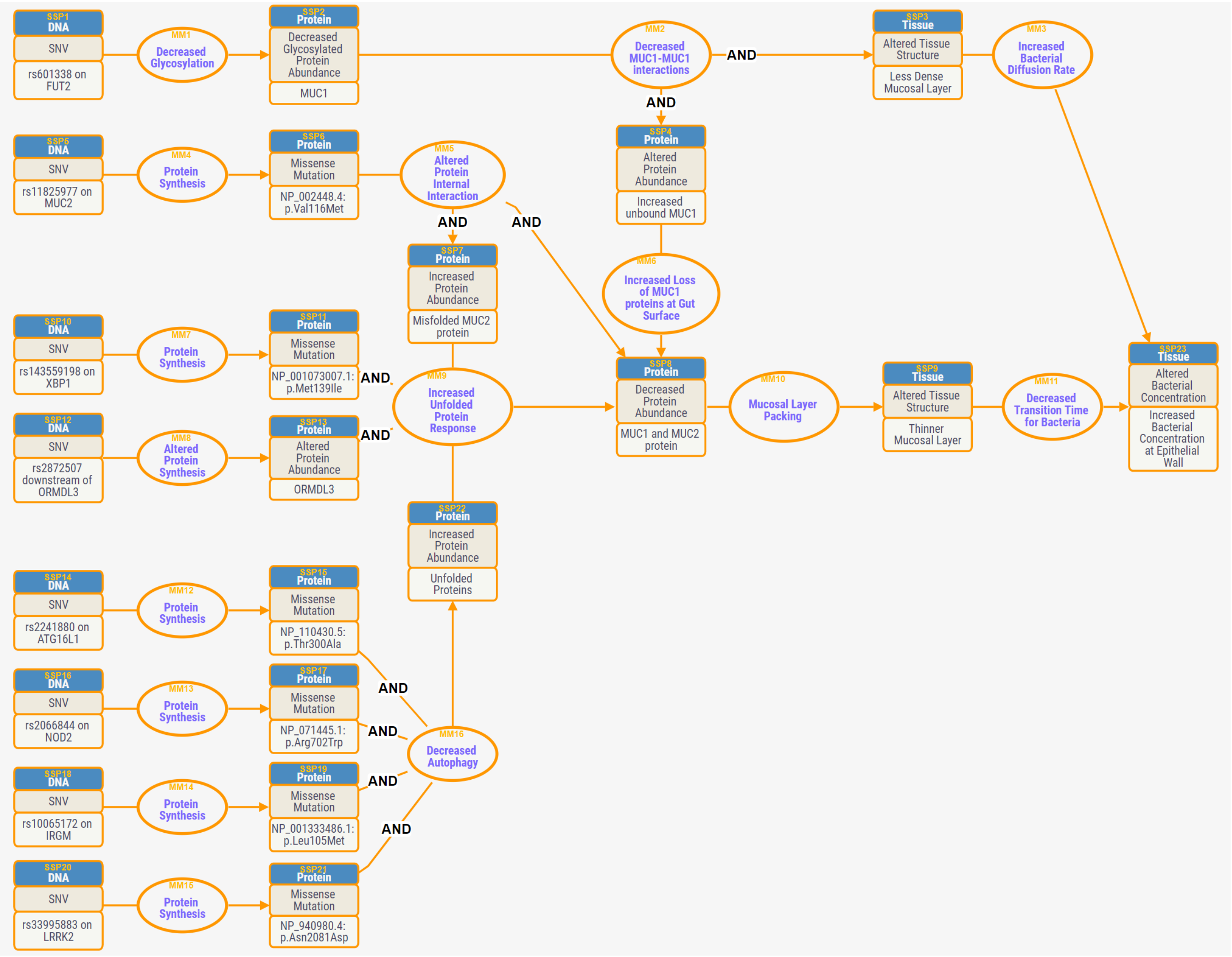


Figure S1: Part of the barrier integrity mechanism graph for Crohn's disease, showing the role of risk variants that affect bacterial penetration through the mucosal layer. SSP=Substate Perturbation and MM=Mechanism Module. An SNP (SSP1) in *FUT2* affects glycosylation (MM1) of MUC1 (Kelly *et al.*, 1995; Mcgovern *et al.*, 2010). The hypoglycosylated state of the MUC1 (SSP2) affects its interaction strength with other mucins (MM2) resulting in a less dense mucosal layer (SSP3) (Hall *et al.*, 2017). Weaker mucin interactions result in more rapid diffusion of bacteria through the mucosal layer (MM3) and faster mucin loss at the gut surface (MM6), one of the three factors contributing to an overall lower abundance of mucin (SSP8). A second factor is a missense SNP in *MUC2 (SSP5)*, the main constituent of gut mucin, resulting in a less stable protein (MM5) (Heazlewood *et al.*, 2008; Moehle *et al.*, 2006). The third factor is the state of the unfolded protein response (UPR) in the mucin producing Goblet cells (MM9). The UPR system reduces protein production in response to the accumulation of excessive misfolded or unfolded protein in the ER (Ma *et al.*, 2017). Because of the normal rapid loss of mucins at the gut interface, Goblet cells are among the most hard-working protein-producing cell types (Gersemann *et al.*, 2009), and so are particularly susceptible to changes in the UPR (Ma *et al.*, 2017). The UPR threshold is influenced by variants in two genes, *XBP1* (Kaser *et al.*, 2008) and *ORMLD3* (Barrett *et al.*, 2008; Moffatt *et al.*, 2007). The extent of misfolded protein is also influenced by the efficiency of autophagy (MM16), involving variants affecting four genes – *NOD2* (Strober *et al.*, 2014), *IRGM* (Chauhan *et al.*, 2016), *LRRK2* (Hui *et al.*, 2018) and *ATG16L1*(Salem *et al.*, 2015).
